# Efficient co-transcriptional splicing enforces rapid microexon definition and inclusion by SRRM4

**DOI:** 10.64898/2026.05.02.722430

**Authors:** Jackson M. Gordon, Joseph Neos Cruz, Karla M. Neugebauer

## Abstract

Alternative splicing expands the coding potential of the genome. Typical human exons are 150 nucleotides long, encoding 50 amino acids. Microexons are only 3-27 nucleotides long; yet they are important regulators of cellular processes in neurons, muscle, and pancreas. In neurons, microexon inclusion is aided by binding of the neuronal splicing factor SRRM4 to flanking upstream 3’ splice sites (3’SSs). Whether this manner of exon definition can be achieved in the timeframe of co-transcriptional splicing is unknown. Here, we employed nascent RNA sequencing to analyze SRRM4-dependent microexons in neuronal cells and found that co-transcriptional microexon splicing is so efficient, the upstream intron is removed before the downstream intron is completely synthesized. This suggests a mechanism for microexon inclusion, whereby co-transcriptional removal of the upstream intron eliminates competition for the microexon’s non-canonical downstream 5’SS. We found that strengthening this 5ʹSS promoted constitutive microexon inclusion independently of SRRM4, indicating that SRRM4 binding alone is a strong stimulator of microexon definition. Thus, SRRM4’s role is to promote rapid splicing of the upstream intron, leaving the microexon’s non-canonical 5’SS as the only option for further splicing. These physiologically significant splicing events thereby require co-transcriptionality to yield neuronal mRNA isoforms.

## INTRODUCTION

Pre-mRNA splicing involves the excision of noncoding intronic sequence and ligation of coding exons in a two-step transesterification reaction^1^. Due to the somewhat degenerate nucleotide determinants that comprise mammalian 5ʹ and 3ʹ splice sites, alternative splicing is widespread. Alternative mRNA isoforms can have vastly different properties and are key players in numerous gene expression programs^2^. Alternative splicing is especially prevalent in the brain where it is utilized by neurons to fine tune the activities, localization, and expression levels of proteins related to neuronal development and synaptic function. Although the average internal exon size in mammals is roughly 150 nucleotides, the most highly conserved class of tissue-specific alternative splicing is a subclass of cassette exons termed microexons, ranging from 3-27 nt^3,4^. These exceptionally small exons are predominantly found in mRNA isoforms expressed in the brain and in genes that are enriched for neuronal function^4^; other microexons generate important mRNA isoforms in striated and cardiac muscle as well as pancreatic islet cells^5,6^. Microexons are most often frame-preserving. Consequently, their inclusion can alter protein-protein interaction networks by introducing new binding surfaces or post-translational modification sites. Mis-splicing of microexons was recently found to be a widespread feature in brain tissue derived from individuals with autism spectrum disorder^7^. Additionally, deletion of microexons in the translation factors *Eif4g1* and *Eif4g3* was sufficient to disrupt normal synaptic function in mouse neurons and led to cognitive deficits in mice^8^. The deletion of a 24-nucleotide microexon in the human cytoplasmic polyadenylation element binding protein CPEB4 also leads to aggregate formation and altered expression of autism-linked genes^9^. These studies highlight the functional importance of maintaining microexon splicing fidelity throughout development.

Despite their clear importance, the reduced size of microexons presents an apparent complication for their recognition by the spliceosome. Splicing in mammals is largely co-transcriptional, with introns often removed rapidly soon after the 3′ splice site is synthesized^10^. For microexons, the 5′ and 3′ splice sites will emerge from the RNA polymerase II (Pol II) exit channel nearly simultaneously. Thus, it is puzzling how microexons are recognized rapidly enough to be spliced into transcripts before competition with the downstream 3′ splice site arises. Several studies have provided insight into how microexons are recognized by the spliceosome. Transcriptomic analyses of post-mortem brain tissue from patients with autism revealed a decrease in expression of the splicing factor SRRM4 (also referred to as nSR100)^7^. A follow-up CRISPR-Cas9 screen confirmed SRRM4’s role in microexon recognition, and identified two other protein factors, SRSF11 and RNPS1 that help promote microexon inclusion^11^. Additionally, inclusion of different subclasses of microexons can be promoted by RBFOX or repressed by PTBP1^4^. More recently, the U1 snRNP subunit PRPF40A was shown to be important for microexon recognition, particularly for smaller microexons^12^. However, these studies have relied on analyzing mature mRNA splice isoforms rather than splicing intermediates. Consequently, little is known regarding co-transcriptional nature of microexon splicing and how the transcriptional environment may impact microexon inclusion levels. Microexon inclusion has been observed in chromatin-associated RNA, indicating microexons can be spliced co-transcriptionally^13^. Delayed splicing of some microexons could create opportunities for regulation by downstream cis-regulatory elements. On the other hand, efficient co-transcriptional splicing of the upstream intron could force inclusion of the microexon before transcription of the downstream intron is complete. Previous work from our lab and others has indicated splicing in mammals is widely co-transcriptional, and introns are often removed in the order they are synthesized^10,14,15^. Co-transcriptional splicing has a high potential to impact alternative exon inclusion due to crosstalk between Pol II, the splicing machinery, and auxiliary splicing factors, and often establishes discrete window of time and space in which splicing can take place^16^. In this study, we determine the extent to which microexons are spliced co-transcriptionally and investigate the co-transcriptional landscape in which microexons are spliced compared to longer constitutive and alternative exons. By applying short- and long-read sequencing methods previously established in our lab^10,17^ to the mouse neuroblastoma Neuro-2a cell line, we detected rapid splicing of SRRM4-dependent microexons, similar to longer exons. Additionally, some SRRM4-independent microexons were spliced later than longer exons, suggesting some microexons may be more susceptible to changes in transcription than others. Overall, our study provides a global analysis of alternative microexon splicing during transcription, including evidence that efficient upstream intron removal can determine splicing outcome.

## RESULTS

### Microexons can be spliced co-transcriptionally in N2a cells

As a model system to study co-transcriptional microexon splicing, we used Neuro-2a (N2a) cells, a robust and rapidly growing cell line capable of expressing neuronal splice isoforms that contain microexons^11,18^. Chromatin-associated RNA was isolated from N2a cells by subcellular fractionation (Figure 1A, Figure S1A), followed by poly(A)+ RNA and rRNA depletion. Samples were then subjected to Illumina short-read RNA sequencing, generating roughly 250 million uniquely mapped reads (Table S1). To broadly assess co-transcriptional splicing in N2a cells, we first utilized the splicing per intron metric (SPI), which is calculated from the ratio of sequencing reads that span a given splice junction to reads that span related exon-intron junctions^19^. Generally strong agreement in SPI values between individual replicates was observed (Figure S1B). The median SPI in our nascent RNA-seq short read dataset was 0.81 (Figure 1B), indicating that 81% of introns are removed co-transcriptionally overall (mean value per intron, three biological replicates).

**Figure 1.**
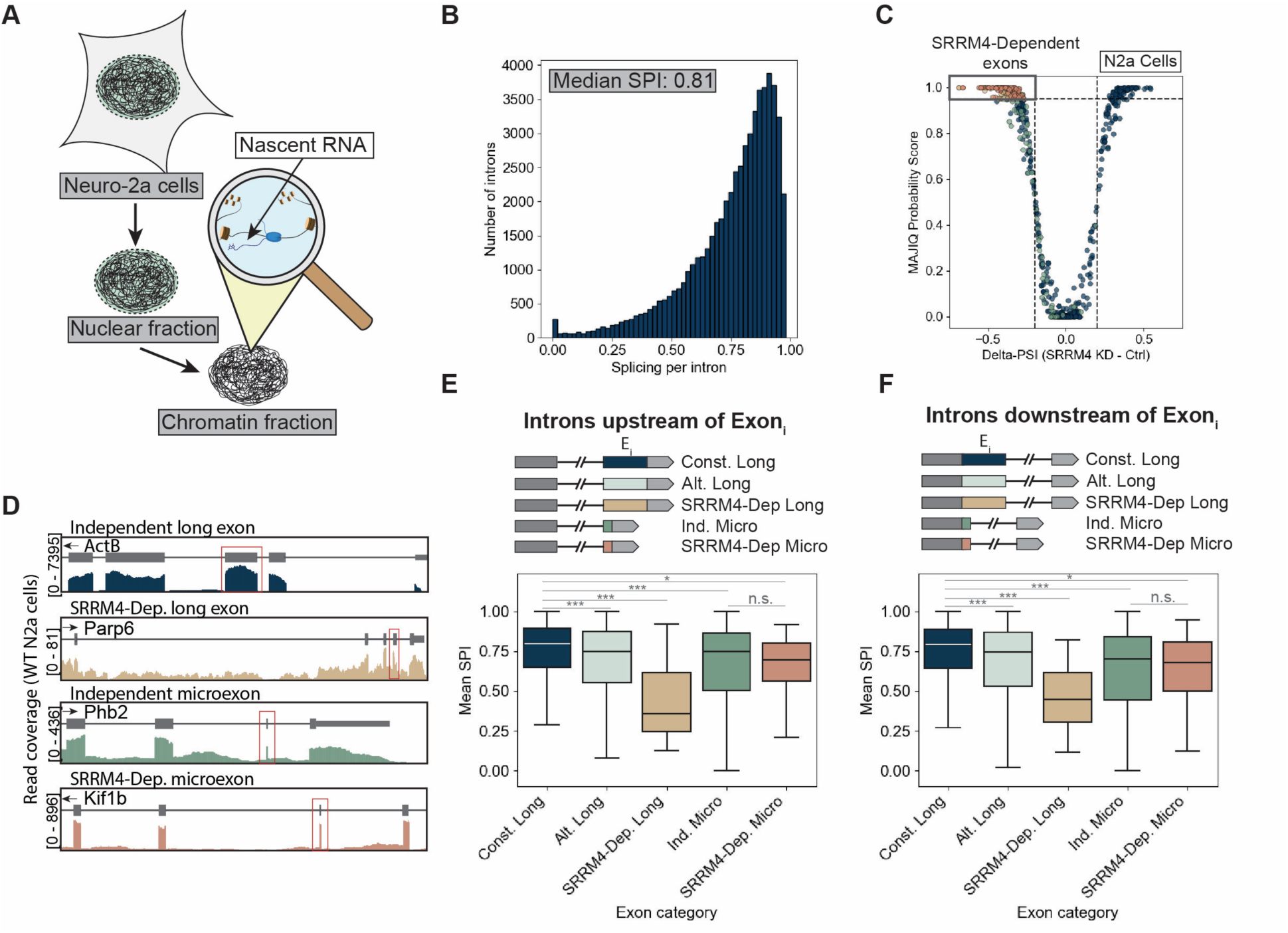
Microexons are efficiently co-transcriptionally spliced. **(A)** Schematic of subcellular fractionation and isolation of nascent RNA. **(B)** Histogram of splicing per intron (SPI) values calculated from short-read sequencing of N2a nascent RNA. **(C)** Volcano plot of cassette exons that are alternatively spliced in N2a cells following siRNA knockdown of SRRM4, RNA-seq data from Raj et al. 2014. MAJIQ was used to quantify alternative splicing events and 95% probability, and 0.2 PSI were used as significance cutoffs. Exons are colored according to length and SRRM4-dependence based on PSI and probability cutoff (Independent long exons: blue; SRRM4-dependent long exons: tan; Independent microexon: green; SRRM4-dependent microexon: red). **(D)** Zoomed in genome browser view of read coverage for *ActB* (Independent long exon), *Parp6* (SRRM4-dependent long exon), *Phb2* (Independent microexon), *Kif1b* (SRRM4-dependent microexon). Read coverage is indicated on y-axes. Exon of interest is highlighted by box. **(E)** Boxplots of mean splicing per intron values for introns upstream of exon of interest (Ei), where Ei represents constitutive long exons (>27 nt) (n=66,038), alternative long exons (n=14,798), SRRM4-dependent long exons (n=23), SRRM4-independent microexons (≤27 nt) (n=391), and SRRM4-dependent microexons (n=24). Statistical significance was determined using a Mann-Whitney U test, *p<0.05,**p<0.005,***p<0.0005. Schematic of introns for which SPI values were calculated is indicated above plots. **(F)** Boxplots of mean splicing per intron values for introns downstream of exon of interest (Ei), where Ei represents constitutive long exons (>27 nt) (n=35,039), alternative long exons (n=10,081), SRRM4-dependent long exons (n=21), SRRM4-independent microexons (≤27 nt) (n=308), and SRRM4-dependent microexons (n=25). Statistical significance was determined using a Mann-Whitney U test, *p<0.05,**p<0.005,***p<0.0005. Schematic of introns for which SPI values were calculated is indicated above plots.

To further categorize alternative exons in N2a cells, we sought to sort exons based on their size and dependence on the known microexon splicing factor SRRM4 for their inclusion. To first classify SRRM4-dependent exons specifically expressed in N2a cells as SRRM4-dependent or independent, we analyzed previously published RNA-seq data from SRRM4 siRNA knockdown N2a cells (from Raj et al., 2014)^18^ using MAJIQ^20^. In total, 229 cassette exons were significantly altered by SRRM4 knockdown (Figure 1C) and skipped exons were disproportionally short (<30 nt) in line with SRRM4’s known role in promoting microexon inclusion (Figure S1C). We utilized the results of this analysis to define discrete categories of alternative exons. Constitutive exons were defined as exons >27 nt that were present in all annotated isoforms of a gene and independent of SRRM4 for their inclusion. Alternative exons were defined as exons >27 nt that were present in at least one but not all isoforms of a gene. SRRM4-dependent long exons were defined as exons >27 nt that were skipped upon SRRM4 KD. Independent microexons were defined as exons ≤27 nt that were independent of SRRM4 for their inclusion. SRRM4-dependent microexons were defined as exons ≤27 nt that were skipped upon SRRM4 KD.

We observed varying coverage across introns flanking SRRM4-independent and SRRM4-dependent exons in our nascent RNA-seq dataset (Figure 1D), prompting us to further explore co-transcriptional splicing patterns in N2a cells. Introns flanking alternative exons generally had lower SPI than those flanking constitutive exons (Figure 1E, Figure 1F), in agreement with previous reports. Introns flanking SRRM4-dependent long exons also had significantly lower SPI than those flanking longer constitutive exons (Figure 1E, Figure 1F). Although introns flanking microexons had slightly lower SPI than longer constitutive exons, the difference was notably smaller than observed for longer SRRM4-dependent exons, indicating microexons may still be spliced rapidly during transcription, while SRRM4 -dependent longer exons might be spliced post-transcriptionally.

### Long read sequencing reveals co-transcriptional coordination of splicing in N2a cells

To more directly define the relationship between splicing kinetics and Pol II position in N2a cells, we prepared long-read sequencing libraries from chromatin-associated RNA using strategies previously devised by the Neugebauer Lab^21^. The 3′-end of each RNA (corresponding to Pol II position) was marked by linker ligation prior to reverse transcription, PCR amplification, and PacBio long-read sequencing. Reads containing poly(A) tails were filtered out prior to mapping. MisER was used to correct for microexon mis-mapping after initial mapping to the *mm39* genome, a common complication with long read mapping algorithms^22^. In total, the filtered library consisted of 532,476 uniquely mapped reads with a median read length of 752 nt, although we observed reads as long as ∼7000 nt (Figure S2A). Our dataset contained 3,449 genes with at least 10 mapped reads in all three biological replicates (Figure S2B) and read coverage was primarily found within genes (Figure S2C).

Our data represent the first chromatin-associated RNA long-read sequencing dataset in a N2a cells. To measure the extent of co-transcriptional splicing in our long-read dataset, we categorized reads as “all spliced”, in which all introns covered in a read were removed, “partially spliced”, in which a fraction of introns in a read were removed, or “all unspliced”, in which no introns had been spliced out (Figure S2D). Partially spliced and all unspliced reads had slightly longer read length distributions compared all spliced reads (Figure S2E). Substantial variation in co-transcriptional splicing between genes could also be seen (Figure 2A). Reads mapping to *Eif4g1* and *Calr* were overwhelmingly all spliced, while reads mapping to *Plxna3* and *Nop56* were predominantly partially spliced or all unspliced respectively. Notably, reads mapping to *Calr* were previously reported to be mostly all unspliced in murine erythroid leukemia cells using an identical library preparation strategy^10^, emphasizing the view that different cell types can have drastically different co-transcriptional splicing patterns. To quantitatively measure co-transcriptional splicing efficiency (CoSE), we used an intron-centric scoring metric previously established in our lab analogous to SPI^10^. The median CoSE in our nascent long-read sequencing dataset was 0.94 (Figure 2B), indicating high levels of co-transcriptional splicing for the nascent transcripts profiled from N2a cells. To ensure our observed splicing efficiency distribution was not overly skewed by PCR amplification during library preparation, we examined how well CoSE scores correlated with SPI from matched short-read sequencing datasets (Figure 2C). Although we did observe a slight bias towards higher splicing efficiency scores in the long-read data, relatively strong correlation could still be seen (Pearson’s r 0.72), indicating our long-read dataset is indeed a reliable resource for probing co-transcriptional splicing dynamics in N2a cells.

**Figure 2.**
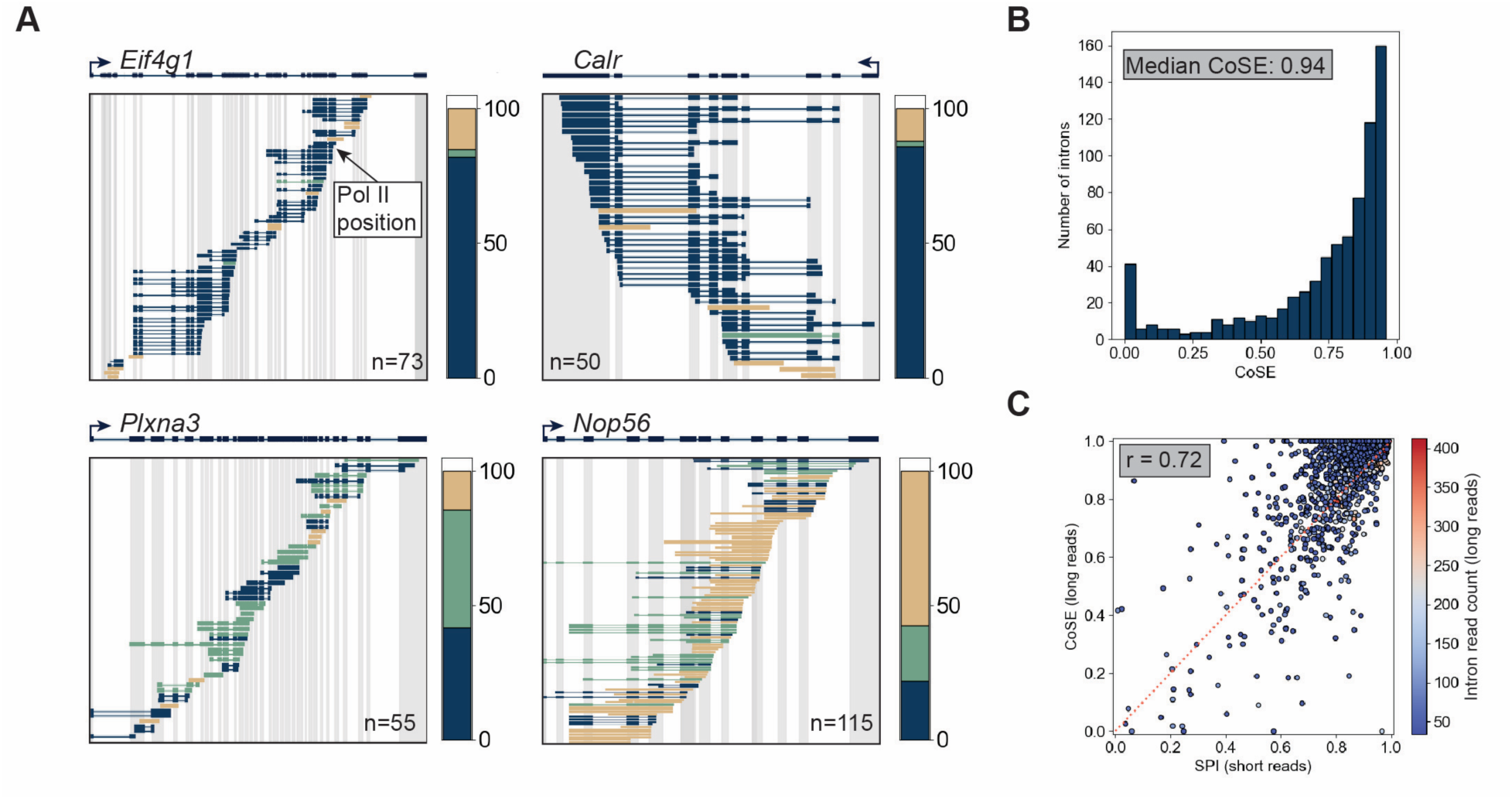
Long-read sequencing of nascent RNA reveals gene-specific co-transcriptional splicing behavior in N2a cells. **(A)** Examples of long reads aligned to *Calr, Eif4g1, Plxna3, and Nop56*, colored by splicing status (All spliced: blue, Partially spliced: green, All unspliced: tan). Percent of reads aligning to each gene in each category is depicted in bar plot to the right of each plot. Number of reads (all replicates combined) is indicated in each plot. Spliced-out introns are indicated by thin lines. **(B)** Histogram of mean co-transcriptional splicing efficiency values (CoSE) calculated for each intron. **(C)** Scatterplot of SPI vs. CoSE calculated for each intron using short and long reads respectively. Pearson’s correlation coefficient was used to assess correlation and is displayed in plot. Color indicates intron long-read coverage for individual introns. Short read and long-read samples were prepared from the same biological replicates of nascent RNA.

### Microexons are rapidly co-transcriptionally spliced

We next wondered if microexons could be spliced rapidly and efficiently with respect to the progression of Pol II (i.e. the distance Pol II has traveled downstream of the microexon), as previously reported for longer exons using similar sequencing methods^10^. Reads containing microexons could be easily visualized in an integrated genomics viewer (Figure S3A). Thus, our dataset confirms some microexons can be spliced co-transcriptionally. First step splicing intermediates, RNAs which have undergone the first transesterification reaction only, were filtered out of the dataset as their 3′-ends, which align exactly at 5′ splice sites, do not represent Pol II position^10^. We next calculated the distance between Pol II and the most proximal upstream splice junction for all reads in the dataset (Figure 3A). The median distance Pol II transcribed before splicing could occur was 176 nt (Figure 3B), similar to previous analyses from our lab in murine erythroid leukemia cells^10^. Interestingly, mean distance Pol II transcribed before an intron is excised and the intron’s splicing efficiency (CoSE) were only slightly negatively correlated indicating that the frequency of co-transcriptional splicing (efficiency) is distinct from the “speed” with which introns are spliced out (Figure 3C). Note that our interpretation of speed is based on Pol II position, and previous analyses have shown that transcription elongation occurs up to 8-fold faster in introns compared to exons^23^; nevertheless, the absence of transcriptional pausing enables us to interpret the distance Pol II has traveled past the microexon as a relative speed of co-transcriptional splicing^10^.

**Figure 3.**
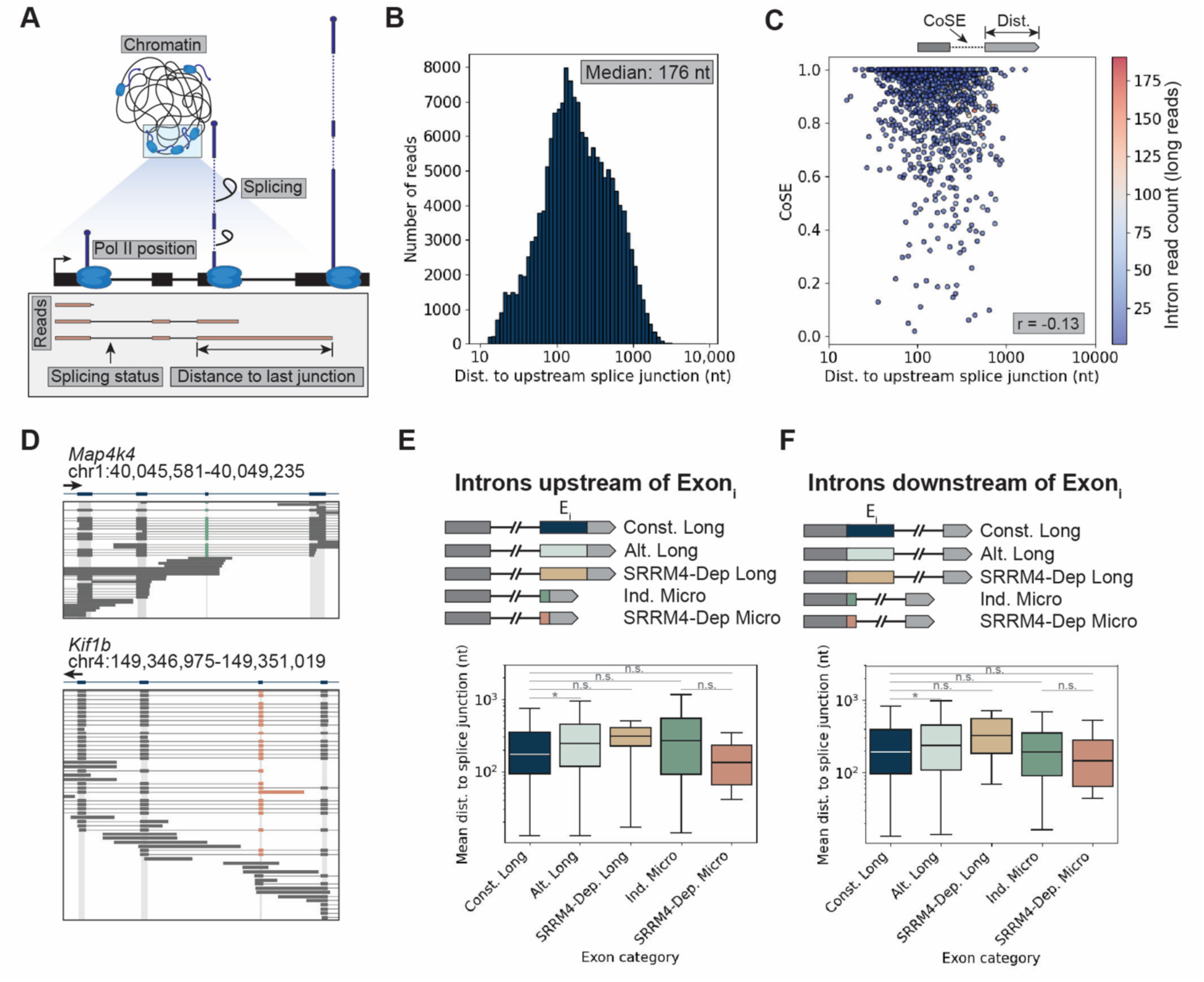
Introns upstream of microexons are rapidly spliced out. **(A)** Schematic depicting how splicing status and Pol II position are derived from long-read sequencing of nascent RNA. **(B)** Histogram of distance in nucleotides between Pol II and last upstream splice junction (e.g. the distance Pol II transcribed into the gene body before splicing can take place). **(C)** Scatterplot of mean distance to upstream splice junction in nucleotides vs. CoSE for each intron. Long-read coverage for each intron is indicated by color. Pearson’s correlation coefficient was used to assess correlation and is displayed in plot. **(D)** Zoomed-in Examples of long reads aligned to *Map4k4,* and *Kif1b.* Exon of interest is colored according to length and dependence on SRRM4 for inclusion (Independent microexon: green, SRRM4-dependent microexon: red). Spliced-out introns are indicated by thin lines. **(E)** Mean distance Pol II transcribes before the intron upstream of an exon of interest (Ei) is excised, where Ei represents constitutive long exons (>27 nt) (n=38,191), alternative long exons (n=10,377), SRRM4-dependent long exons (n=7), SRRM4-independent microexons (≤27 nt) (n=148), and SRRM4-dependent microexons (n=4). Statistical significance was determined using a Mann-Whitney U test, *p<0.05. **(F)** Mean distance Pol II transcribes before the intron downstream of an exon of interest (Ei) is excised, where Ei represents constitutive long exons (>27 nt) (n=19.595), alternative long exons (n=4,553), SRRM4-dependent long exons (n=8), SRRM4-independent microexons (≤27 nt) (n=111), and SRRM4-dependent microexons (n=10). Statistical significance was determined using a Mann-Whitney U test, *p<0.05.

We next separated long reads based on the identity of their last spliced-in exon (referred to here as Ei), in which Ei represents constitutive long exons, alternative long exons, SRRM4-dependent long exons, independent microexons, and SRRM4-dependent microexons. Reads containing individual examples of both microexon categories could be seen in our long-read dataset (Figure 3D, Figure S3B). Pol II transcribed significantly further into the gene body before introns upstream or downstream of long alternative exons could be spliced compared to introns upstream of long constitutive exons (Figure 3E, Figure 3F). Interestingly, introns upstream of SRRM4-independent or dependent microexons were spliced just as rapidly with respect to Pol II position as those upstream of longer constitutive exons. A similar trend was observed for introns downstream of microexons. To determine if the splicing reaction itself is altered around microexons, we analyzed first step splicing intermediates (Figure S3C). However, we observed very few introns with reads corresponding to first step splicing intermediates, indicating the splicing cycle itself also proceeds efficiently around both longer exons and microexons in N2a cells (Figure S3D). Overall, our results indicate that SRRM4-dependent microexons may be rapidly spliced before competition with downstream splice sites arises.

### Targeted long-read sequencing reveals variable splicing of microexons between genes

Our initial observations indicate that globally, SRRM4-dependent microexons are spliced in a relatively short spatial window of ongoing transcription elongation, similar to longer constitutive exons. However, due to limitations in sequencing depth, we were unable to reliably measure differences in co-transcriptional splicing of individual microexons. To improve sequencing depth and resolution of Pol II positions for microexon-containing genes, long-read sequencing libraries were generated using a targeted amplification approach. Matched cDNA samples were reamplified using gene-specific forward primers for microexon containing genes *Ank2*, *Kif1b*, *Mon2*, and *Clasp2*, each of which contained well-characterized, SRRM4-dependent microexons^4^. To ensure adequate capture of the microexon-containing region of the gene, forward amplification primers were designed complementary to the exon immediately upstream of the microexon (Figure 4A, Figure S4A). In parallel, forward primers in the first exon were also used to generate separate libraries. Reverse primers were complementary to the 3′ adapter sequence, capturing Pol II position, consistent with our global long-read sequencing library preparation. In total, 24 separate libraries were prepared (four genes, two forward primer sets each, three biological replicates) and pooled for PacBio long-read sequencing. The final library consisted of 222,390 uniquely mapped targeted reads that aligned to target genes (all datasets combined). The median read lengths for reads amplified with a first-exon complementary primer was 1,353 nt (Figure 4B, Figure S4B-E). The median read lengths for reads amplified with an upstream-exon complementary primer was 1,585 nt (Figure 4B, Figure S4F-I).

**Figure 4.**
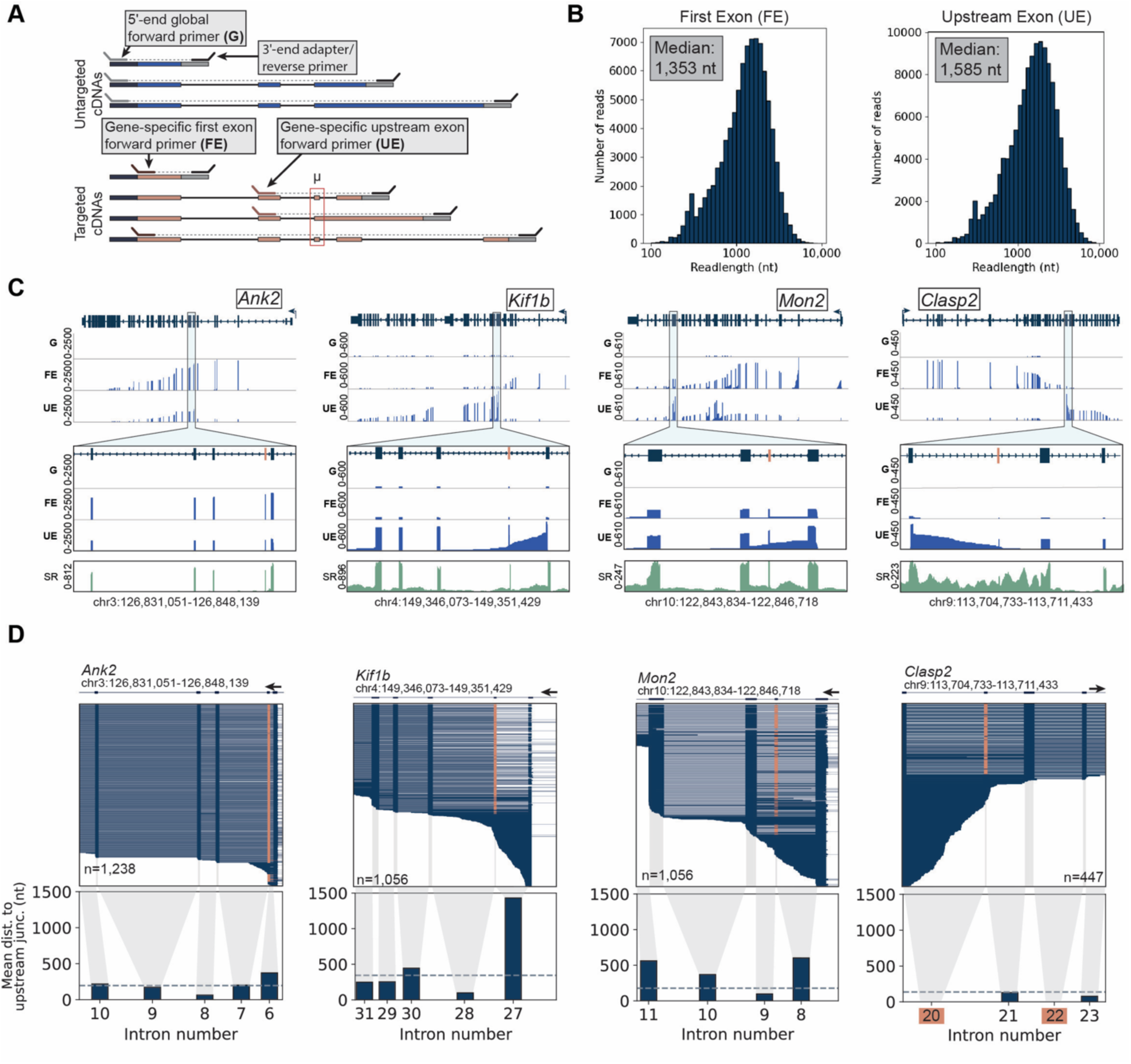
Co-transcriptional splicing varies among individual microexons. **(A)** Schematic of targeted PCR amplification strategy for generating gene-specific long-read sequencing libraries. **(B)** Histogram of read lengths for three biological replicates (all replicates combined) of PacBio long-read sequencing datasets prepared from N2a nascent RNA and amplified with gene-specfic forward primers in first exons. **(C)** Read coverage plots from long-read sequencing libraries prepared by targeted PCR amplification of *Ank2*, *Kif1b*, *Mon2*, and *Clasp2*. Gene diagram is displayed at the top of each plot. Long-read coverage of for libraries prepared using global amplification (G), amplification with a forward primer in the first exon of each gene of interest (FE), and amplification with a forward primer in the exon upstream of the microexon of interest (UE) are displayed in the middle with zoom-ins indicated by each window. Short-read (SR) coverage from global nascent RNA-seq dataset is displayed in green at the bottom of each plot for comparison. **(D)** Zoomed-in examples of long reads aligned to *Ank2, Kif1b, Mon2,* and *Clasp2.* Constituitive exons are colored in blue. Microexons are colored in red. Spliced-out introns are indicated by thin lines (top). Mean distance between Pol II position and upstream splice junctions for introns in corresponding reads plots (bottom). Introns with no data are boxed in red. Median distance to upstream splice junctions for the entire gene is indicated by dashed gray line.

Compared to our original global long-read sequencing dataset, we observed a substantial increase in reads spanning for microexon-containing regions with sufficient sequencing depth to analyze co-transcriptional splicing for individual exons (Figure 4C). We also found that libraries prepared with forward primers complementary to the exon upstream of microexons of interest (UE) better matched short-read sequencing coverage than those prepared with forward primers complementary to first exons (FE), possibly due to combined effects of limited reverse transcriptase processivity and PCR amplification bias (Figure 4C). UE 5′-ends also had single predominant peaks in coverage and 3′-ends were mostly downstream of the amplified exon, while FE 5′- and 3′-ends were slightly more dispersed (Figure S4J-S4M). We therefore decided to use UE datasets for downstream analyses. Variation between the four amplified genes in Pol II position and upstream alternative exon inclusion could be visualized easily in individual reads (Figure 4D). We next applied the previously described metrics to analyze co-transcriptional splicing in the targeted long-read sequencing datasets. The intron upstream of the *Ank2* microexon was rapidly excised, similar to other introns in the gene, and the intron immediately upstream of the *Mon2* microexon was excised at only slightly longer Pol II distance compared to introns further downstream (Figure 4D). Interestingly, Pol II transcribed over 1kb before the intron upstream of the *Kif1b* microexon could be excised and the *Clasp2* microexon was only detected after splicing of downstream introns. These results highlight substantial variation in co-transcriptional splicing dynamics around microexons, possibly due to differences in their absolute dependence on SRRM4 and other auxiliary splicing factors.

### Microexon splice site strength impacts SRRM4 dependence but not co-transcriptional splicing efficiency

The observed variability in co-transcriptional microexon splicing kinetics suggests that although SRRM4 may promote rapid co-transcriptional splicing of some microexons, others may be rely on the presence of additional cis- or trans- regulatory factors. For example, SRRM4-dependent microexons have previously been reported to have stronger than average 5′ splice sites and weaker than average 3′ splice sites (based on consensus sequences)^7^. SRRM4 has also been shown to bind introns upstream of SRRM4-dependent microexons, in between the polypyrimidine tract and 3′ splice site, and relies on the presence of a UGC sequence motif for binding^11^. We therefore sought to investigate the relationship between splice site strength and co-transcriptional microexon inclusion. To first determine whether splice site strength may differentiate rapidly spliced SRRM4-dependent microexons from independent microexons, we used a maximum entropy model (MaxEnt) to score 5′ and 3′ splice sites for exons covered in our long-read sequencing dataset (Figure 5A)^24^. Independent and SRRM4-dependent microexons both had slightly stronger 5′ splice sites, in agreement with previous reports, though not significant between exons only in our dataset. Longer SRRM4-dependent exons had weaker 5′ splice sites compared to longer exons (constitutive and alternative combined), possibly explaining the observed lower splicing efficiency of flanking introns. Both longer exons and microexons that were SRRM4-dependent had weaker 3′ splice sites compared to exons that are spliced independent of SRRM4. SRRM4-independent microexons had relatively strong 3′ splice sites (Figure S5A), suggesting combined strong 5′ and 3′ splice sites may be sufficient to promote constitutive microexon inclusion in N2a cells.

**Figure 5.**
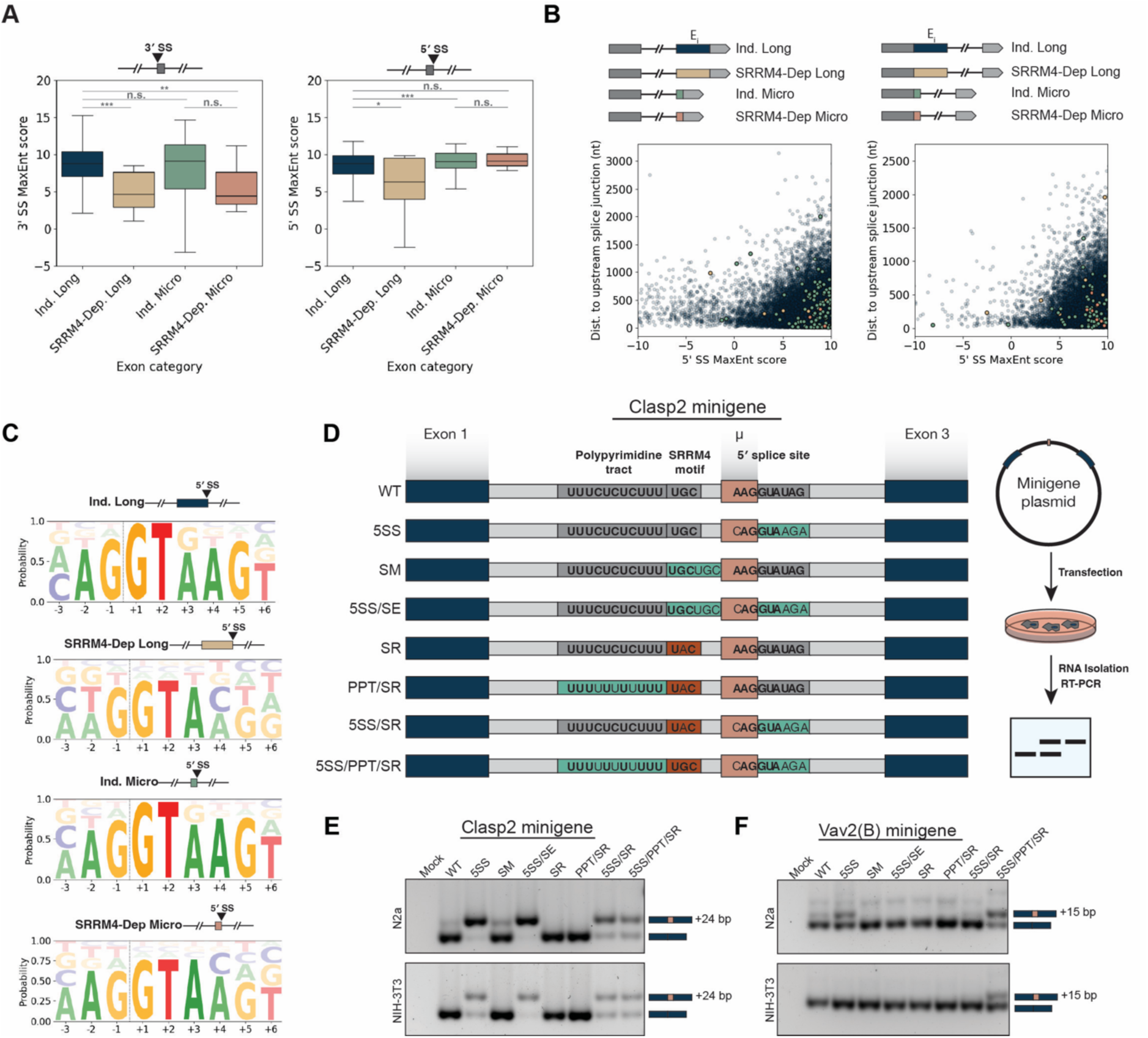
Consensus 5’ splice sites promote microexon inclusion in select SRRM4-dependent microexons. **(A)** Maximum entropy scores for 3’ splice sites (left) and 5’ splice sites (right) for exons with significant coverage in long-read sequencing dataset. **(B)** Maximum entropy scores for 5’ splice sites vs. distance between Pol II and upstream splice junctions for introns upstream of exons of interest (left) or downstream of exons of interest (right). **(C)** Logo plots of 5′ splice sites of independent long exons, independent microexons, SRRM4-dependent long exons, and SRRM4-dependent microexons generated using exons found in global long-read sequencing dataset. Junction positions are indicated by horizontal lines. **(D)** *Clasp2*-based minigene constructs. Teal regions indicate mutations expected to enhance strength of motif. Red regions indicate mutations expected to hamper strength of motif. Bolded nucleotides indicate changes to consensus splice sites or polypyrimidine tracts. Blue boxes represent exons, while gray boxes represent introns. Salmon box indicates 24-nucleotide microexon. Schematic of minigene transfection and RT-PCR is shown to the left of minigene constructs. **(E)** RT-PCR gel for Clasp2-based minigene constructs. Upper bands represent microexon-included isoform while lower bands represent microexon-skipped isoform. **(F)** RT-PCR gel for Vav2(B)-based minigene constructs. Upper bands represent microexon-included isoform while lower bands represent microexon-skipped isoform.

We next investigated whether splice site strength may impact how far Pol II must transcribe before an exon is spliced into a nascent RNA. Interestingly, there was little correlation between distance to the upstream splice junction and 5′ splice site MaxEnt scores (Figure 5B). Thus, stronger splice sites strength may promote spliceosome assembly but do not necessarily promote more rapid co-transcriptional splicing. We also note that although microexon splice sites are sometimes stronger than longer independent exons, they are not often true consensus 5′ splice sites (complementary to U1 snRNA) (Figure 5C). We wondered how splice sites ultimately *combine* with other cis-regulatory elements to ultimately determine the frequency of inclusion for SRRM4-dependent microexons. We therefore decided to further investigate the impact of combinatorial sequence elements on microexon inclusion in N2a cells. To do so, we turned to a minigene reporter system, which enables higher throughput mutational screening than CRISPR-Cas9-mediated mutagenesis or overexpression systems.

Minigenes were cloned into a pcDNA5 vector under regulation by a cytomegalovirus promoter for robust expression in mammalian cells. Initially, a minigene based on the endogenous *Clasp2* sequence was generated (Figure 5D). Iterations harboring various combinations of stronger (more pyrimidine-rich) polypyrimidine tracts (PPTs), SRRM4-binding motif mutations, and 5′ splice site sequences were then prepared to probe the combined effect of these elements on microexon splicing. We first focused on a 24-nucleotide long microexon in *Clasp2*, which our gene-specific sequencing data indicated was poorly co-transcriptionally spliced. The wild-type *Clasp2* minigene produced only slight inclusion of the 24 nt microexon (Figure 5E). Interestingly, by altering the 5′ splice site to exactly compliment U1 snRNA, near constitutive inclusion of the microexon was observed. Extension of the SRRM4-bindding motif also bolstered microexon inclusion to a lesser extent. As expected, mutating the UGC to UAC in the SRRM4-binding motif abrogated microexon inclusion. Surprisingly, microexon inclusion could not be rescued in the presence of this mutation by increasing the pyrimidine richness in the PPT. Rather, a stronger 5′ splice site was required to rescue some amount of microexon inclusion in the UAC point mutant. To rule out possible contributions by SRRM4 to microexon inclusion in any of the above mutational contexts, minigenes were transfected into the mouse fibroblast NIH-3T3 cell line, as these cells are not known to express SRRM4. Interestingly, similar patterns of alternative splicing were observed, with strong promotion of microexon inclusion in mutants with strong 5′ splice sites (Figure 5E).

To examine whether stronger 5′ splice sites promoted microexon inclusion in other gene contexts, the same mutations were prepared and tested using the endogenous sequence of *Vav2(B)*, which contains a 15 nt microexon (Figure S5B). In N2a cells, the presence of a stronger 5′ splice site *Vav2(B)* mutant boosted levels of microexon inclusion compared to wild-type, consistent with initial results from the *Clasp2* minigene (Figure 5F). However, in the presumed absence of SRRM4 binding (in the UAC point mutant or in NIH-3T3 cells), a stronger polypyrimidine tract and stronger 5′ splice site were both required to promote microexon inclusion. These results confirm that by tuning cis-regulatory elements in and surrounding microexons, SRRM4-dependent microexons can be made independent of SRRM4. However, the exact contributions of the 5′ splice and PPT to microexon inclusion appear to be highly context dependent. A recent study indicated strong 5′ splice sites can compensate for loss of SRRM4 in promoting microexon inclusion in human cells^25^. Our results support an underappreciated role for 5′ splice sites in microexon splicing more broadly and highlight possible differences in cis-regulatory elements between SRRM4-dependent and SRRM4-independent microexons.

## DISCUSSION

In this study, we provide a co-transcriptional perspective of SRRM4-dependent microexon recognition by the splicing machinery. Our custom nascent RNA-seq datasets yield measurements of the efficiency and order of intron removal as well as the distance Pol II travels before two exons are ligated together. Combined with our minigene approach that identifies the importance of the SRRM4 binding site relative to the splice sites, we propose the working model schematized in Figure 6. Because upstream and downstream intron removal is highly co-transcriptional and occurs when Pol II is less than 200 nt downstream of the respective 3′ SS, the upstream intron is predominantly removed before the downstream intron, making microexon inclusion inevitable. Subsequent removal of the downstream intron is similarly fast and efficient. Therefore, 3′ splice site recognition upstream of the microexon, which is dependent on SRRM4 in neurons, must be very rapid. SRRM4 likely associates with its cognate binding site concurrently with its synthesis by Pol II. This inference is reinforced by the demonstration that artificially strengthening the downstream 5′ SS can render the microexon 100% included and independent of SRRM4, as expected for an exon definition mechanism. Therefore, many natural microexons must be defined by the rapid co-transcriptional binding of SRRM4, enabling substantial tissue-specific microexon inclusion.

**Figure 6.**
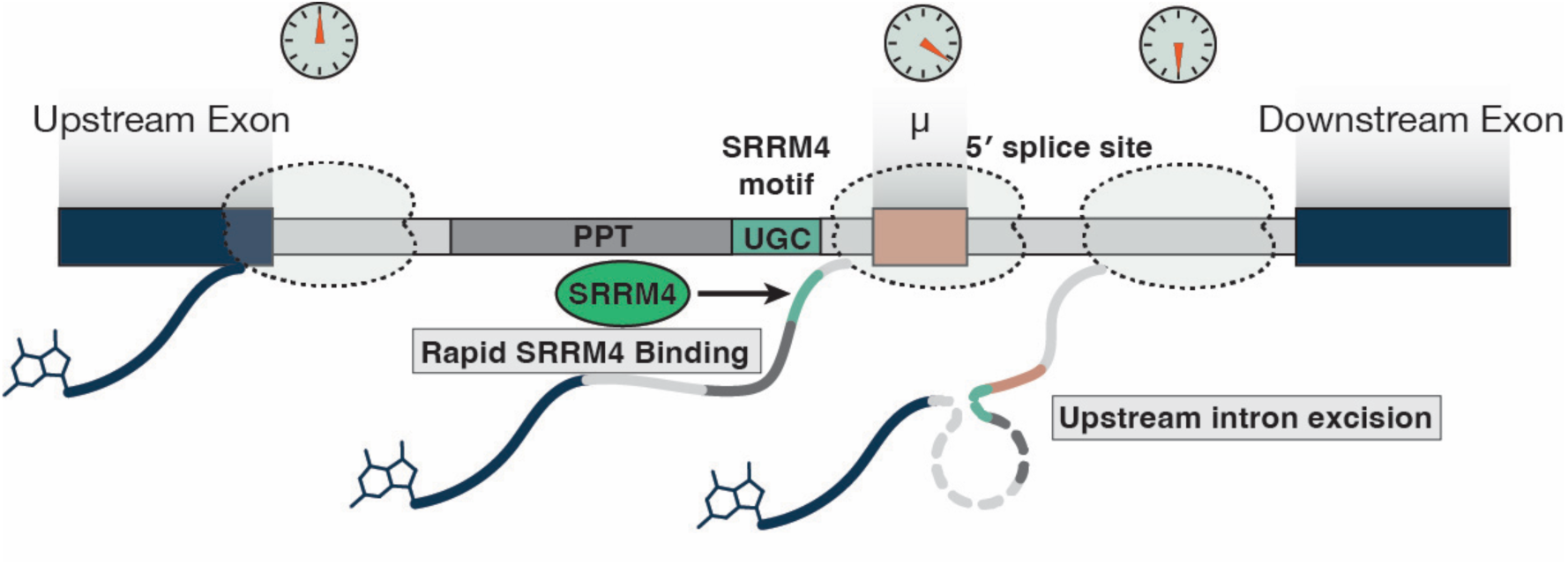
Working model of co-transcriptional splicing of SRRM4-dependent microexons. SRRM4 binds upstream introns rapidly after synthesis of its binding motif by Pol II. Rapid binding of SRRM4 promotes excision of the intron upstream of the microexon, ensuring the mature mRNA isoform contains the microexon.

While the focus of this study is the challenge of understanding how microexons can be co-transcriptionally defined and included in neurons, our datasets have also enabled a global-scale description of co-transcriptional splicing in a neuronal cell line for the first time. Work from our lab and others has indicated that splicing in mammals is largely co-transcriptional and distinct patterns of intron excision have been observed between genes^10,15^. Short read sequencing of chromatin-associated nascent RNA isolated from N2a cells revealed that 81% of introns are spliced out co-transcriptionally. Both short-read and long-read sequencing detected widespread rapid and efficient co-transcriptional splicing in these cells. Interestingly, cell type-specific differences in co-transcriptional splicing could be readily observed by comparing long reads with published studies. For example, reads mapping to the gene *Calr* were efficiently spliced co-transcriptionally in N2a cells compared to their very inefficient splicing, with intron retention across most whole transcripts, in murine erythroid leukemia cells^10^. Other genes like *Nop56* are inefficiently spliced across multiple introns in neurons, providing a counterexample. Finally, even though efficient co-transcriptional splicing is widespread in N2a cells, intron- and gene-specific behaviors span a wide range of splicing efficiencies, highlighting the potentially powerful role delayed splicing may play in determining gene output, since unspliced transcripts are retained in the nucleus and often targeted for RNA decay^26,27^.

Recent work has demonstrated potential biochemical bases for the coupling of transcription and splicing. U1 snRNP has been shown to interact directly with Pol II *in vitro*^28^. Additionally, the Pol II subunit RPB9 was shown to mediate co-transcriptional splicing by interacting with U2AF1, promoting early 3′ splice site recognition^29^. In the co-transcriptional landscape of splicing in N2a cells, microexons were still able to be spliced into nascent pre-mRNAs as rapidly as longer constitutively spliced exons, no matter whether the exon was dependent or independent of SRRM4 for its inclusion. SRRM4-independent microexons had stronger 5′ and 3′ splice sites which may account for their rapid co-transcriptional splicing. In contrast, for SRRM4-dependent microexons, which are frequently spliced into neuronal transcripts due to tissue-specific expression of SRRM4, SRRM4 appears to mediate early formation of an exon definition complex that allows the exon to be spliced into nascent pre-mRNAs before synthesis of the competing downstream 3′ splice site is complete (see Figure 6). Whether a specific mechanism of SRRM4 recruitment, such as tracking with elongating Pol II or its concentration within a transcription and splicing condensate, is needed to explain the rapid definition of microexons remains to be established. For that matter, how tracking of U1 snRNP, U2AF, or other splicing factors with Pol II contribute to microexon splicing is also unknown.

Local transcription rates have been shown to affect inclusion rates of alternative exons^30^. The slow Pol II mutant *Polr2a* R749H was embryonic lethal in homozygous mice^31^, likely at least partially due to defective splicing that is essential for normal development. In embryonic stem cells heterozygous for this mutation, an increased frequency of longer cassette exons was observed compared to wild type, indicating slow Pol II may favor exon inclusion. Conversely, microexons were more frequently skipped in the same mutant. This suggests differing regulatory mechanisms at the co-transcriptional level for longer cassette exons and microexons, many of which are dependent on SRRM4. Microexon splicing may have evolved from intronic sequence with the emergence of SRRM4 homologues^3^. Indeed, Pol II is reported to transcribe faster over introns than exons ^23^. Thus, it is possible that SRRM4 has evolved to favor the environment surrounding faster Pol II to slower Pol II. Additional work probing co-transcriptional microexon splicing in slow Pol II mutants will be required to dissect the optimal spatial relationship between Pol II and microexon-promoting factors.

Previously established models of microexon definition indicate microexons are first recognized by a complex containing SRRM4, SRSF11, and RNPS1 which recruits U1 and U2 snRNPs for early spliceosome assembly^4^. PRP40A, a component of the U1 snRNP, was also recently shown to promote microexon splicing in N2a cells^12^. Thus, by ensuring strong interaction between microexon 5′ splice sites and U1 snRNP, the requirement for PRP40A appears to overcome the challenge presented by weaker polypyrimidine tracts or lack of exonic splicing enhancer motifs. These observations point to the interesting question of whether microexons characteristic of other tissues, such as muscle or pancreatic islets, are also regulated co-transcriptionally by other tissue-specific splicing factors. Because microexon mis-splicing is a frequent occurrence in human disease and developmental disorder, inducing autism^4^, mechanistic insights into the dynamics of microexon definition and splicing during transcription elongation contribute to the overall goal of therapeutically modulating microexon inclusion in neurons as well as other tissues.

## METHODS

### Cell Lines and Cell Culture

Neuro-2a (N2a) cells^32^ were cultivated in Dulbecco’s modified Eagle’s medium GLutaMAX (DMEM) (Gibco) supplemented with 10% fetal bovine serum (Gibco) and 1% penicillin/streptomycin (Gibco). Cells were cultured at 37°C, 5% CO2.

### Western Blotting

Western blots were carried out as previously described^33^ using antibodies specific for the following proteins/epitopes: GAPDH (abcam ab9485), RNA Polymerase II (4H8) (Santa Cruz sc-47701), U1-70K (CB7) (hybridoma supernatant), Histone H3 (abcam ab1791). Total protein was stained with Ponceau S (Thermo Scientific) as recommended by the manufacturer. Western blots were developed with ECL Western Blotting Detection Kit (Cytiva Amersham) and visualized using a Bio-Rad GelDoc imaging system.

### Biochemical Fractionation and Nascent RNA Isolation

Biochemical fractionation and nascent RNA isolation was performed essentially as described in^21^. N2a cells were harvested by rinsing in ice cold PBS followed by centrifugation at 270 x g for 10 min. Cell pellets were washed twice in ice cold PBS. Three biological replicates were prepared (roughly 20 million cells per replicate) from cells grown on three separate days. Cell pellets were resuspended 250 µL of cell lysis buffer containing 10 mM Tris-HCl pH 7.5, 0.1% NP-40, 75 mM NaCl, 40 U/mL SUPERase.IN (Thermo Fischer Scientific, AM2694), 1x cOmplete protease inhibitor mix (Sigma) and incubated on ice for 7 minutes to dissociate the cell membrane while leaving nuclei intact. Lysate was then layered on top of 500 µL of buffer containing cell lysis buffer with 24% (w/v) sucrose and centrifuged for 10 minutes at 6,000 x g, 4°C (Eppendorf 5427 R). The cytoplasmic fraction was then aspirated off nuclei. Nuclei were washed with 500 µL of ice-cold PBS/1 mM EDTA before gently resuspending in 100 µL of resuspension buffer containing 20 mM Tris-HCl pH 8.0, 75 mM NaCl, 0.5 mM EDTA, 0.85 mM DTT, 50% glycerol, 40 U/mL SUPERase.IN, 1x cOmplete protease inhibitor mix. 100 µL of nuclear lysis buffer containing 20 mM HEPES pH 7.5, 1 mM DTT, 7.5 mM MgCl2, 0.2 mM DTDA, 300 mM NaCl, 1 M Urea, 1% NP-40, 40 U/mL SUPERase.IN, 1x cOmplete protease inhibitor mix was added and samples vortexed for 5 seconds followed by 3-minute incubation on ice to lyse nuclei. Chromatin was pelleted by centrifugation for 2 minutes at 6,000 x g, 4°C. Nucleoplasm was removed from chromatin and chromatin was rinsed with 500 µL of PBS/1mM EDTA. Chromatin was then resuspended in 100 µL of PBS and 300 µL of Trizol. At each fractionation step, 50 µL aliquots were removed for analysis by western blot as described above. Pol II 4H8, U1-70K, and GAPDH were used as chromatin, nucleoplasmic, and cytoplasmic markers respectively (Figure S1A). Chromatin samples in Trizol were incubated in a thermomixer at 50°C for 10 minutes with shaking at 1,400 rmp. Chromatin-associated RNA was then isolated by Trizol extraction according to the manufacturer. Further RNA cleanup was carried out using the RNeasy Mini kit (Qiagen) according to the manufacturer using on-membrane DNA digestion with DNase I (Qiagen). Poly(A)+ RNA was depleted from RNA samples using the DynaBeads mRNA Direct Micro Purification kit (Thermo Fischer Scientific) according to the manufacturer. Finally, ribosomal RNA was depleted from samples using the RiboMinus Eukaryote kit (Thermo Fischer Scientific) according to the manufacturer.

### Short-Read RNA-seq Library Preparation

Nascent RNA was prepared from N2a cells as described above and used to prepare RNA-seq libraries using the KAPA mRNA HyperPrep Kit (Kapa Biosystems). The RNA-seq library was sequenced on the Illumina Novaseq 6000 platform generating ∼30 million paired-end reads per sample.

### RNA-seq Data Analysis

Paired-end read mates were generated in. fastq format. Illumina adapters were trimmed using fastp^34^. Reads were mapped to the mouse GRCm39 (mm39) genome using STAR^35^ with settings --runMode alignReads --outSAMtype BAM SortedByCoordinate --outFilterMismatchNmax 2 --alignIntronMin 20 --alignIntronMax 1000000 --alignMatesGapMax 1000000. Mapped files were indexed using samtools index^36^. SPLICE-q was used to calculate splicing per intron values^37^. For reproducibility, a Snakemake^38^ pipeline that carries out the processing steps outlined above was generated. Downstream analysis and plotting were carried out using custom python scripts.

### Transcriptome-Wide Long-Read Sequencing Library Preparation

Nascent RNA was prepared from N2a cells as described above. Transcriptome-wide sequencing library preparations were performed essentially as described in^21^. A custom DNA adapter (Table S2) was ligated to the 3′ ends of nascent RNAs using the T4 RNA ligase kit (NEB) according to the manufacturer. Adapter-ligated RNAs were reversed transcribed using the SMARTer PCR cDNA Synthesis kit (Takara) with the following modifications: the CDS Primer IIA was replaced with a primer complementary to the custom adapter sequence with Primer IIA sequence overhang. The resulting cDNA was PCR amplified to 18 cycles using the Advantage 2 PCR (Takara) according to the manufacturer using the SMARTer PCR cDNA Synthesis kit Primer IIA (complementary to both ends of the cDNA). DNA purification was carried out using AMPure XP beads (Beckman Coulter) and pooled at equimolar ratios. The PacBio SMRTbell prep kit was used for final library preparation according to the manufacturer (Pacific Biosciences) and sequenced on a PacBio Sequel II Long-Read Sequencer (Table S1).

### Gene-Specific Long-Read Sequencing Library Preparation

For improved sequencing depth of genes of interest, targeted PCR amplification of previously prepared cDNA libraries was carried out. Four genes (*Ank2*, *Mon2*, *Kif1b*, *Clasp2*) were selected for targeted amplification based on the following criteria: (1) genes contain annotated microexons, (2) genes have evidence of microexon inclusion in the transcriptome-wide nascent RNA sequencing dataset, and (3) microexons within each gene have been suggested to alter function of the corresponding protein^4^. Forward primers were complementary to regions either in Exon 1 or the first upstream exons of the microexon for each gene. Reverse primers were complementary to barcode sequences in the 3′ ends of cDNA (part of the ligated adapter sequence). cDNA was PCR amplified to 16 cycles using the Advantage 2 PCR (Takara) according to the manufacturer. DNA purification was carried out using AMPure XP beads (Beckman Coulter) and pooled at equimolar ratios. The PacBio SMRTbell prep kit was used for final library preparation according to the manufacturer (Pacific Biosciences) and sequenced on a PacBio Sequel II Long-Read Sequencer.

### Long-Read Sequencing and Data Analysis

HiFi circular consensus sequencing (CCS) reads were generated in. fastq format. For downstream removal of PCR duplicates, a custom python script was used to append the 5-nucleotide UMI sequence to the header of each read. Reads were then mapped to the mouse GRCm39 genome (mm39) using minimap2^39^ with settings -x splice:hq –secondary no -t 12 -a -u f –seed 14. Samtools^36^ was used to convert .sam files to .bam format and sort reads according to genomic coordinates. MisER was then used to correct for misaligned microexons^22^. UMICollapse^40^ was used to remove PCR duplicates and bedtools^41^ intersect was used to filter out small fragments that only mapped to intronic regions. Finally, custom python scripts were used to filter polyadenylated reads and first step splicing intermediates (reads with 3′ ends at 5′ splice site positions). For reproducibility, a Snakemake^38^ pipeline that carries out the processing steps outlined above was generated. Calculation of distance to splice junctions was carried out using custom python scripts. Maximum Entropy scores were calculated according to Yeo and Burge, 2004^24^. Plots were generated using matplotlib and seaborn python packages.

### Minigene Design and Strategic Considerations

Microexon-containing minigenes were designed based on the endogenous gene architecture and sequences of mouse Clasp2 and Vav2(B). Wild-type constructs contained the respective microexon sequence for each gene of interest and the immediate flanking upstream and downstream intron and exon. Minigene sequences were cloned into a pcDNA5 vector^11^. Candidate genes were initially selected based on the presence of an annotated microexon that is dependent on the spicing factor SRRM4 for their inclusion^4^. Additionally, candidates produced both microexon-included and skipped isoforms in N2a cells. Mutant constructs were prepared with combinations of the following mutations: Polypyrimidine tracts in the intron upstream of the microexon were strengthened by conversion to poly(U). The UGC motif between the polypyrimidine tract and microexon 3′ splice site was weakened by mutation to UAC and enhanced by extension to UGCUGC. Finally, the microexon 5′ splice site was strengthened by conversion to the consensus sequence CAG/GUAAGU.

### Minigene Cloning and Mutagenesis

Minigene sequences for Claps2 and Vav2(B) were PCR amplified (Phusion High Fidelity DNA Polymerase, NEB, see Table for primer sequences) from N2a genomic DNA isolated using the Quick-DNA Miniprep kit (Zymo Research). PCR products were gel purified with a 1% agarose gel and Qiagen DNA Gel Extraction kit according to the manufacturer. Amplicons were cloned into pcDNA5/FRT/TO vector (Invitrogen) using the In-Fusion HD Cloning kit (Takara) according to the manufacturer. NotI restriction enzyme (NEB) was used to linearize the plasmid prior to In-Fusion cloning. Full constructs were transformed and isolated from Stellar Competent Cells (Takara) and plasmids were sequenced in full (Plasmidsaurus). Mutagenesis was carried out using the Quick Change Lighting Site-Directed Mutagenesis kit (Agilent) according to the manufacturer (see Table for mutagenesis primer sequences. All mutations were confirmed using Sanger sequencing (Quintara Bio) and whole plasmid sequencing (Plasmidsaurus or Quintara).

### Minigene Transfection and Assessment by RT-PCR

Minigenes were transiently transfected into N2a or NIH-3T3 cells using the Lipofectamine 3000 reagent (ThermoFisher) according to the manufacturer 24 hours after seeding 100,000 - 150,000 cells/well in 12-well plates. 1 µg of plasmid was used for transfection. Cells were then incubated at 37°C for 48 hours. Total RNA was isolated by TRIzol extraction according to the manufacturer and reverse transcribed to cDNA using SuperScript III Reverse Transcriptase (ThermoFisher) and random hexamer primer mix. PCR amplification of the minigene from cDNA was performed with Phusion High Fidelity Polymerase (NEB) and custom primers in which the forward primer was complementary to a region between the promoter and first exon and reverse primer complementary to a region in the last exon. PCR products were separated on a 2% agarose TBE gel (MetaPhor agarose, Lonza Biosciences) for 5 hours, 120V. Gels were stained with GelStar Nucleic Acid Gel Stain (Lonza Biosciences) and bands corresponding to microexon skipped and included minigenes were visualized using a Bio-Rad GelDoc imaging system.

## RESOURCE AVAILABILITY

### Lead Contact

Further information and requests for resources and reagents should be directed to and will be fulfilled by the lead contact, Karla Neugebauer

## Data and Code Availability

All short and long-read RNA sequencing data generated in this study have been uploaded to the Gene Expression Omnibus (GEO) and are publicly available under the accession number (GSE3095990). All original code has been deposited on GitHub (https://github.com/NeugebauerLab/N2a_LRS.git) and is publicly available.

## ACKNOWLEDGEMENTS

We thank Benjamin Blencowe for the gift of wild-type Neuro-2a cells. We are grateful to members of the Neugebauer lab for helpful discussions. This work was supported by the National Institutes of Health (R01 GM112766 to K.M.N and F31NS129248 to J.M.G.). Its contents are solely the responsibility of the authors and do not necessarily represent the official views of the NIH. We thank the Yale Center for Genomic Analysis for providing the sequencers and assisting in library preparation. Research reported in this publication was supported by the National Institute of General Medical Sciences of the National Institutes of Health under Award Number 1S10OD030363-01A1.

## AUTHOR CONTRIBUTIONS

J.M.G and K.M.N designed this study. J.M.G. and J.N.C. performed subcellular fractionations and RNA sequencing library preparations. J.N.C. constructed minigene plasmids and carried out site-directed mutagenesis. J.N.C. and J.M.G. assessed splicing products following transient transfection of minigene plasmids. J.M.G. performed all sequencing data analysis. J.M.G. and K.M.N. wrote the manuscript.

