## Supplemental files for "Efficient co-transcriptional splicing enforces rapid microexon definition and inclusion by SRRM4"

**SUPPLEMENTAL FIGURES and TABLES**

**Supplemental Figure 1.** Fractionation quality assessment and determination of SRRM4-dependent microexons.

**Supplemental Figure 2.** Library characteristics of PacBio long-read sequencing dataset prepared from N2a nascent RNA.

**Supplemental Figure 3.** Co-transcriptional splicing of exons of interest, assessed by long-read sequencing.

**Supplemental Figure 4.** Library characteristics of PacBio long-read sequencing dataset prpared from N2a nascent RNA with targeted amplifcation of genes of interest.

**Supplemental Figure 5.** 3′ splice site sequence characteristics for SRRM4-dependent and independent exons.

**Supplemental Table S1:** Mapping statistics of RNAseq data sets analyzed in this study.

**Supplemental Table S2:** Oligonucleotides used in this study.

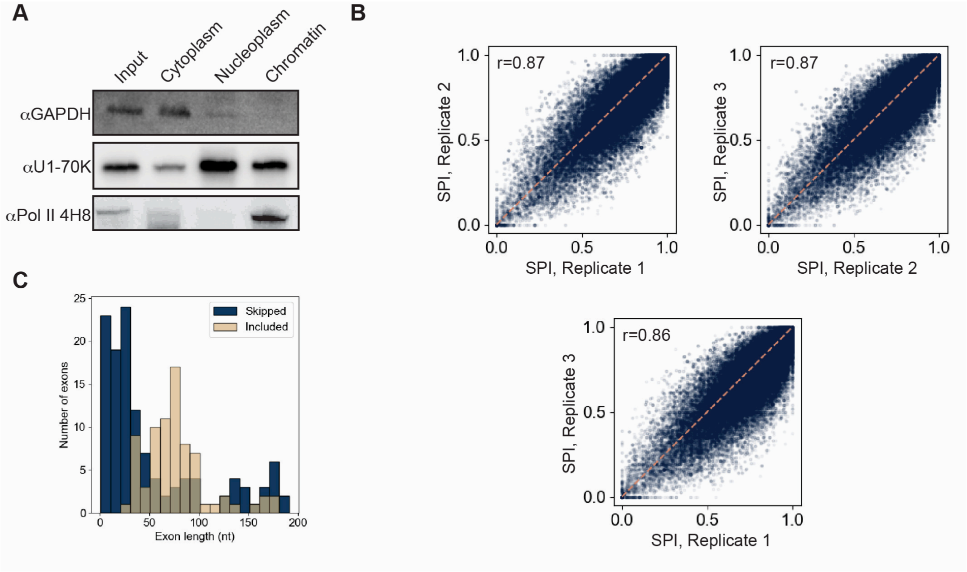


**Figure S1. Fractionation quality assessment and determination of SRRM4-dependent microexons.**

**(A)** Anti-GAPDH, anti-U1-70K, and anti-Pol II 4H8 western blots of input, cytoplasm, nucleoplasm, and chromatin fractions from N2a cells. **(B)** Splicing per intron values (SPI) for individual introns from Replicate 1 vs Replicate 2 (top left), Replicate 2 vs. Replicate 3 (top right), and Replicate 1 vs. Replicate 3 (bottom). Pearsons's correlation coefficient is displayed at top left for each comparison. **(C)** Histogram of exon length of significant skipped and included exons in SRRM4 knockdown compared to control.

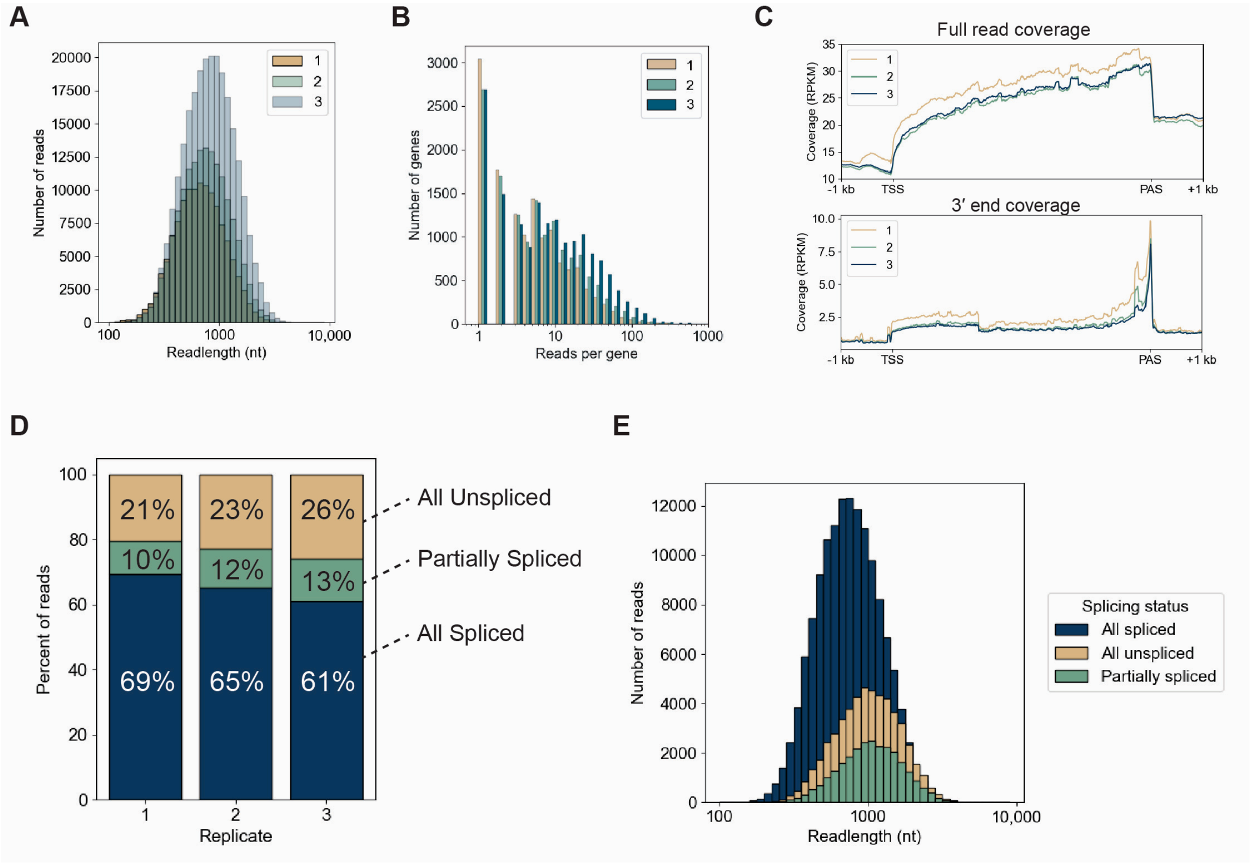


**Figure S2. Library characteristics of PacBio long-read sequencing dataset prepared from N2a nascent RNA.**
**(A)** Histogram of read lengths for three biological replicates of PacBio long-read sequencing datasets prepared from N2a nascent RNA. **(B)** Histogram of read coverage per gene for long-read sequencing dataset, separated by biological replicate. **(C)** Metaplots of coverage for full reads (top) and 3′ ends (bottom), separated by biological replicate. **(D)** Percent of reads in long-read sequencing dataset categorized as “All spliced” (blue), “Partially spliced” (green), and “All unspliced” (tan), separated by biological replicates. **(E)** Histograms of read length distributions separated by splicing status (All spliced, Partially spliced, or All unspliced); biological replicates combined.

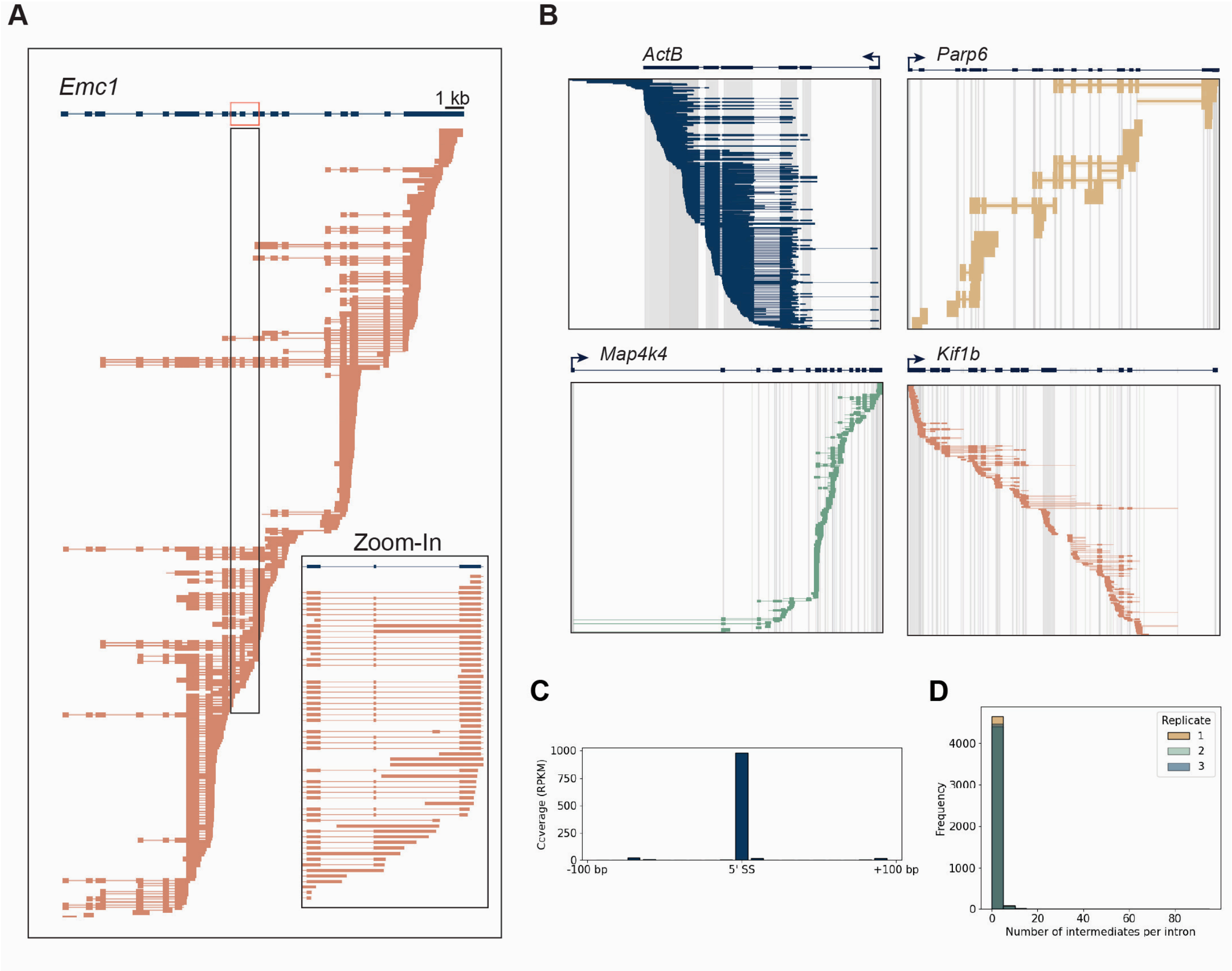


**Figure S3. Co-transcriptional splicing of exons of interest, assessed by long-read sequencing.**

**(A)** Example of long reads from nascent RNA long read sequencing library, prepared from N2a cells. Reads shown are aligned to *Emc1* reference gene (top) and the box indicates reads zoomed in at the microexon-containing segment. Reads displayed are from 3 combined biological replicates from WT N2a cells. **(B)** Full coverage of reads aligned to *ActB, Parp6, Map4k4*, and *Kif1b*, related to **Figure 3D**. Blue reads (ActB) contain only constitutive exons. Tan reads (*Parp6*) contain at least one SRRM4-depdenent long exon. Green reads (*Map4k4*) contain only constitutive exons. Tan reads (*Parp6*) contain at least one SRRM4-depdenent long exon. Green reads (*Map4k4*) contain at least one SRRM4-independent microexon. Salmon reads (*Parp6*) contain at least one SRRM4-depdenent microexon. **(C)** Metaplot of coverage for 5′ ends from reads corresponding to first-step splicing intermediates. **(D)** Histogram of the number of splicing intermediate reads per intron, separated by replicate.

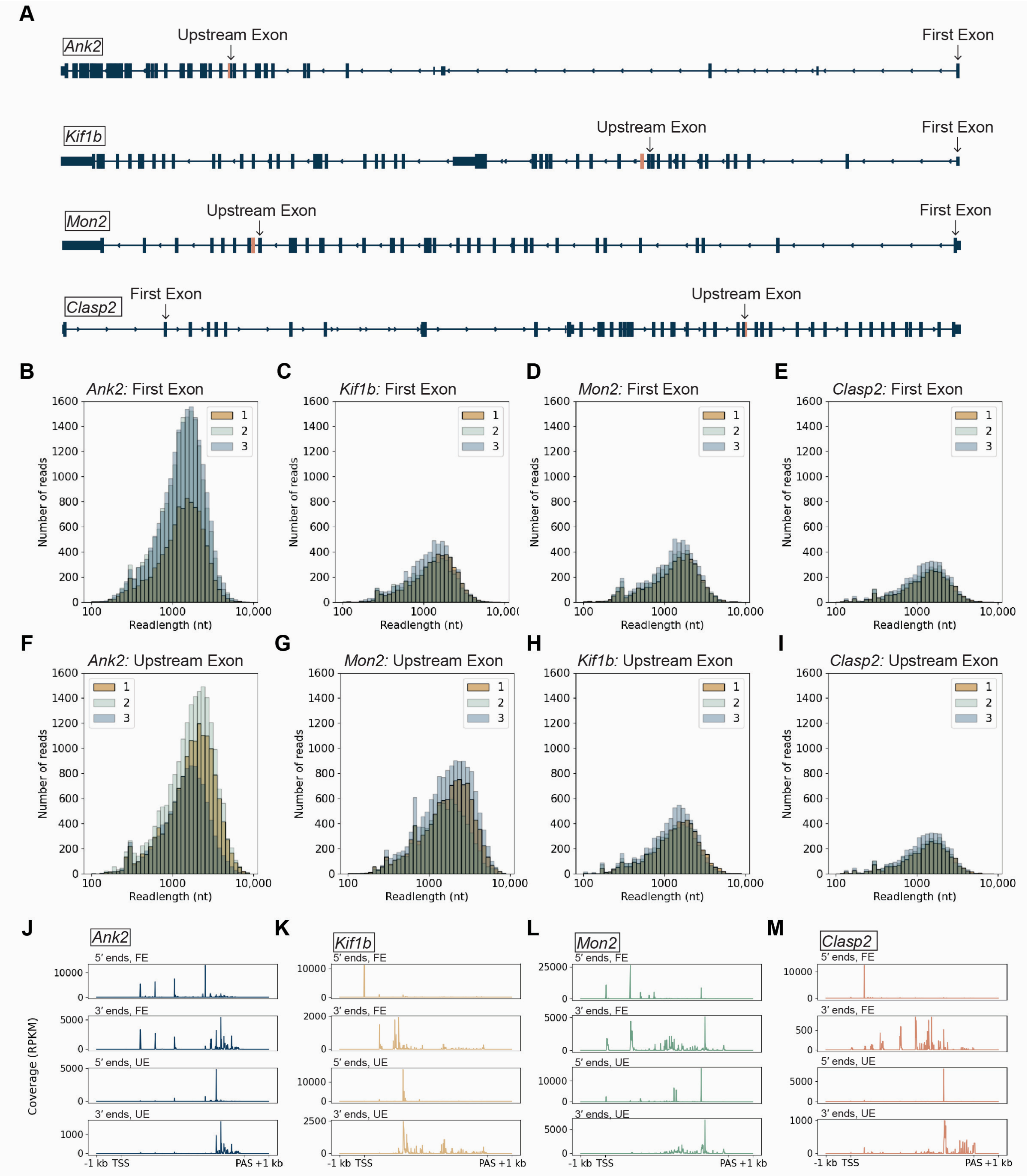


**Figure S4. Library characteristics of PacBio long-read sequencing dataset prpared from N2a nascent RNA with targeted amplifcation of genes of interest.**

**(A)** Forward locations for targeted amplification of *Ank2, Kif1b, Mon2,* and *Clasp2*. **(B-I)** Histograms of read lengths for three biological replicates of PacBio long-read sequencing datasets prepared from N2a nascent RNA with targeted PCR amplification. **(J-M)** Metaplots of coverage for 5'-ends and 3'-ends with first exon (FE) and upstream exon (UP) forward primers for *Ank2* **(J)**, *Kif1b* **(K)**, *Mon2* **(L)**, and *Clasp2* **(M)**.

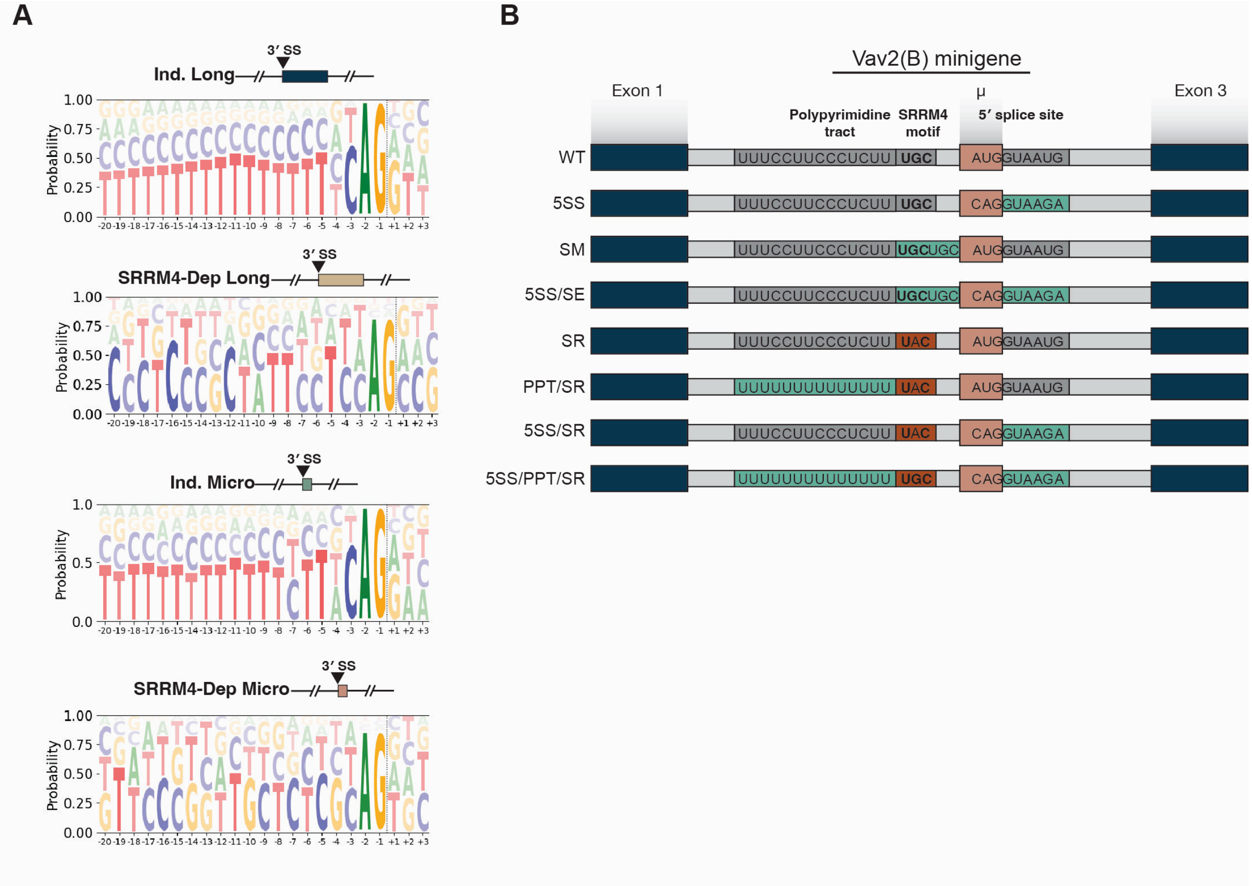


**Figure S5. 3′ splice site sequence characteristics for SRRM4-dependent and independent exons.**

**(A)** Logo plots of 3′ splice sites for independent long exons, independent microexons, SRRM4-dependent long exons, and SRRM4-dependent microexons generated using exons found in global long-read sequencing dataset. Junction positions are indicated by horizontal lines. **(B)** *Vav2(B)*-based minigene constructs. Teal regions indicate mutations expected to enhance strength of motif. Red regions indicate mutations expected to hamper strength of motif. Bolded nucleotides indicate changes to consensus splice sites or polypyrimidine tracts. Blue boxes represent exons, while gray boxes represent introns. Salmon box indicates 15-nucleotide microexon.

**Table S1:** Mapping statistics of RNAseq data sets analyzed in this study.

| Sample | Replicate | Platform | Uniquely mapped reads (mm39) |
|---|---|---|---|
| N2a, Global | 1 | Illumina NovaSeq | 77859267 |
| N2a, Global | 2 | Illumina NovaSeq | 75832998 |
| N2a, Global | 3 | Illumina NovaSeq | 99120126 |
| N2a, Global | 1 | PacBio Sequel II | 915786 |
| N2a, Global | 2 | PacBio Sequel II | 1319505 |
| N2a, Global | 3 | PacBio Sequel II | 2175640 |
| N2a, Ank2, First exon | 1 | PacBio Sequel II | 106712 |
| N2a, Ank2, First exon | 3 | PacBio Sequel II | 150841 |
| N2a, Ank2, First exon | 3 | PacBio Sequel II | 148168 |
| N2a, Ank2, Upstream exon | 1 | PacBio Sequel II | 60419 |
| N2a, Ank2, Upstream exon | 2 | PacBio Sequel II | 137400 |
| N2a, Ank2, Upstream exon | 3 | PacBio Sequel II | 38984 |
| N2a, Mon2, First exon | 1 | PacBio Sequel II | 45817 |
| N2a, Mon2, First exon | 2 | PacBio Sequel II | 20805 |
| N2a, Mon2, First exon | 3 | PacBio Sequel II | 42221 |
| N2a, Mon2, Upstream exon | 1 | PacBio Sequel II | 50219 |
| N2a, Mon2, Upstream exon | 2 | PacBio Sequel II | 37393 |
| N2a, Mon2, Upstream exon | 3 | PacBio Sequel II | 75718 |
| N2a, Kif1b, First exon | 1 | PacBio Sequel II | 15075 |
| N2a, Kif1b, First exon | 2 | PacBio Sequel II | 15811 |
| N2a, Kif1b, First exon | 3 | PacBio Sequel II | 31077 |
| N2a, Kif1b, Upstream exon | 1 | PacBio Sequel II | 30256 |
| N2a, Kif1b, Upstream exon | 2 | PacBio Sequel II | 15003 |
| N2a, Kif1b, Upstream exon | 3 | PacBio Sequel II | 47042 |
| N2a, Clasp2, First exon | 1 | PacBio Sequel II | 15367 |
| N2a, Clasp2, First exon | 2 | PacBio Sequel II | 13817 |
| N2a, Clasp2, First exon | 3 | PacBio Sequel II | 18687 |
| N2a, Clasp2, Upstream exon | 1 | PacBio Sequel II | 37219 |
| N2a, Clasp2, Upstream exon | 2 | PacBio Sequel II | 71366 |
| N2a, Clasp2, Upstream exon | 3 | PacBio Sequel II | 89159 |

### Table S2: Oligonucleotides used in this study

| Description | Sequence |
|---|---|
| Long reads 3′ end adapter | /5rApp/NNNNNCTGTAGGCACCATCAAT/3ddC/ |
| Library prep RT primer, Replicate 1 | AAGCAGTGGTATCAACGCAGAGTACCACATATCAGAGTGCGGATTGATGGTGCCTACAG |
| Library prep RT primer, Replicate 2 | AAGCAGTGGTATCAACGCAGAGTACACACACAGACTGTGAGGATTGATGGTGCCTACAG |
| Library prep RT primer, Replicate 3 | AAGCAGTGGTATCAACGCAGAGTACACACATCTCGTGAGAGGATTGATGGTGCCTACAG |
| Targeted library prep, Ank2, first exon primer | ATGTTTTTTACTCCTCTGGCTGTGGCAC |
| Targeted library prep, Ank2, upstream exon primer | TTAACTAATCGCTGAGAGAAGCTGCTGC |
| Targeted library prep, Mon2, first exon primer | CTGGCATGTGTTGGACCAGAAAACATGCAG |
| Targeted library prep, Mon2, upstream exon primer | GGAAATTGCAAGAAGAAGTTCCCACCTGTC |
| Targeted library prep, Kif1b, first exon primer | ATATACAATATCAATAAGTCCCTTACAACTTTGGGC |
| Targeted library prep, Kif1b, upstream exon primer | AATCTTCTCGGCGTCTTCTGACTCGG |
| Targeted library prep, Clasp2, first exon primer | TTTCGAGATATGGGATCAGCCAGTCCAGC |
| Targeted library prep, Clasp2, upstream exon primer | AATTACGGTGGGCTCGAGCAACTACC |
| Targeted library prep, reverse primer | AAGCAGTGGTATCAACGCAGAGTAC |
| Minigene universal reverse primer | AATTCTGCAGATATCCAGCACAGTG |
| Clasp2 minigene forward amplification primer | ATGATTCATACCTTCTTCGAGCAGG |
| Vav2(B) minigene forward amplification primer | TGAATCAGGTAGAAGCCATAGGACC |
